# pH-Dependent Transcriptional Profile Changes in Iron-Deficient *Arabidopsis* Roots

**DOI:** 10.1101/2020.05.21.109777

**Authors:** Huei-Hsuan Tsai, Wolfgang Schmidt

**Affiliations:** Institute of Plant and Microbial Biology, Academia Sinica, Taipei, 11529, Taiwan; Biotechnology Center, National Chung-Hsing University, Taichung, 40227, Taiwan; Genome and Systems Biology Degree Program, College of Life Science, National Taiwan University, Taipei, 10617, Taiwan

**Keywords:** ambient pH, coumarins, iron deficiency, iron uptake, RNA-seq, transcriptome, alkaline soil

## Abstract

Iron is an essential element for plants and abundantly present in most mineral soils. The mobility of iron is, however, dependent on the redox potential and hydrogen activity (pH) of the soil, factors that may limit its availability to plants in particular at alkaline pHs. Iron deficiency triggers pronounced changes in the transcriptional profile of plants, inducing processes that aid in the acquisition, uptake, and translocation of iron. How ambient pH impact the transcriptional iron deficiency response has not yet been elucidated in detail. Here, we provide an RNA-seq data set that catalogs global gene expression changes of iron-deficient plants grown at either optimal (5.5) or high (7.0) pH. A suite of 857 genes changed significantly and more than twofold in expression; only 54 genes of this suite were also differentially expressed between iron-deficient and iron-sufficient plants grown at pH 5.5. Among the high pH-responsive genes, 186 were earlier shown to be responsive to short-term transfer to low pH, 91 genes of this subset were anti-directionally regulated by high and low pH. The latter subset contained genes involved in cell wall organization, auxin homeostasis, and potential hubs of yet undefined signaling circuits. Growing iron-deficient plants at high pH also modulated the transcriptional iron deficiency response observed at pH 5.5 by compromising the enzymatic reduction of ferric chelates and favoring the production of iron-mobilizing coumarins. It is concluded that ambient pH is an important determinant of global gene expression which tunes iron acquisition to the prevailing edaphic conditions.

## INTRODUCTION

Soil pH, *i.e.* the dynamic equilibrium of H^+^ activity between the soil solution and the negatively charged solid phase, is an important edaphic factor that dictates the availability of mineral nutrients, affects the composition of the microbiome, and determines the composition of plant communities through alterations in the availability of mineral nutrients in the soil (Ellenberg, 1978). Iron is highly abundant in most soils, but the low mobility of oxidized iron compounds often limits its phyto-availability. In aerated soils, the solubility of iron decreases by a factor of 1,000 for each unit increase in pH between 4 and 9, severely restricting the supply of iron at circumneutral or alkaline conditions (Lindsay, 1979).

In *Arabidopsis* and other non-grass species, iron starvation triggers a sophisticatedly regulated response comprising processes which increase the solubilization of recalcitrant soil iron pools, including the acidification of the rhizosphere, secretion of iron-mobilizing compounds, and reductive splitting of ferric chelates with subsequent release and uptake of Fe^2+^ (Kobayashi et al., 2019). Soil pH has a pronounced effect on the acquisition of iron. In acidic soils, high manganese solubility can interfere with iron uptake and may cause secondary iron deficiency (Marschner, 1995). Alkaline conditions, on the other hand, not only restrict the availability of iron, but also compromise the enzymatic reduction of ferric chelates by the plasma membrane-bound reductase FRO2, a central part of the iron uptake mechanism (Susin et al., 1996). Restricted mobilization and limited uptake of iron constitute main factors for excluding so-called calcifuge (‘chalk-fleeing’) plants from habitats with alkaline soil reaction (Tyler, 1996).

While the transcriptional response of *Arabidopsis* to iron starvation has been well explored (e.g. Thimm et al., 2001; Schmidt and Buckhout, 2011; Kim et al., 2019), most studies were conducted at slightly acidic pH which is optimal for growth, but leaves the influence of alkaline pH on global gene expression profiles undefined. Knowledge on how ambient pH, in particular neutral or alkaline conditions, affects the transcriptional landscape of iron-deficient plants may aid in understanding the bottlenecks of gene regulation in plants that are not well adapted to soils with limited iron availability. Here, we provide an RNA-seq-based inventory of genome-wide gene expression of roots from iron-deficient *Arabidopsis* plants grown either at optimal, slightly acidic (5.5), or high (7.0) pH. To mimic the restricted availability of iron at high pH, we provided an iron source of low solubility to plants grown at neutral pH. Alkaline soil reaction aggravates iron deficiency symptoms, but the effect of pH *per se* has not yet been clearly distinguished from the response to a lack of iron at a hydrogen activity that is optimal for growth. We believe that the data set provided here allows for gaining valuable insights into the underlying causes of restricted iron uptake limiting growth in natural or agronomical ecosystems with high pH, and sets the stage for follow up experimentation aimed at identifying novel players involved in the adaptation of plants to the prevailing hydrogen activity.

## MATERIALS AND METHODS

Seeds of the *Arabidopsis* (*Arabidopsis thaliana*) accession Col-0 were obtained from the Arabidopsis Biological Resource Center (Ohio State University). Plants were grown under sterile conditions in a growth chamber on agar-based media as described by Tsai et al. (2018). The growth medium comprised 5 mM KNO_3_, 2 mM MgSO_4_, 2 mM Ca(NO_3_)_2_, 2.5 mM KH_2_PO_4_, 70 μM H_3_BO_3_, 14 μM MnCl_2_, 1 μM ZnSO_4_, 0.5 μM CuSO_4_, 0.01 μM CoCl_2_, and 0.2 μM Na_2_MoO_4_, supplemented with 1.5% (w/v) sucrose, and solidified with 0.4% Gelrite pure (Kelco). For iron-deficient optimal-pH media, 100 μM ferrozine and 1 g/L MES were added, and the pH was adjusted to 5.5 with KOH. For iron-deficient high-pH media, 40 μm FeCl_3_ and 1 g/L MOPS were added, and the pH was adjusted to 7.0 with KOH. Plants were grown on media for 14 d.

For RNA-seq analysis, total RNA was isolated from roots of 14-d-old plants grown under iron-deficient conditions either at optimal (5.5) or high (7.0) pH using the RNeasy Plant Mini Kit (Qiagen). Libraries for RNA-seq were prepared by using the Illumina TruSeq RNA library Prep Kit (RS-122-2001, Illumina) according to the manufacturer’s protocol. Briefly, 4 μg of total RNA per sample were used for library construction. PolyA RNA was captured by oligodT beads and fragmented when eluted. cDNA was synthesized from fragmented RNA using reverse transcriptase (SuperScript III, Cat. No. 18080-093, Invitrogen) and random primers. Reactions were cleaned up with Agencourt AMPure XP beads (Beckman Coulter Genomics). Libraries were end-repaired, adenylated at the 3’ end, ligated with adapters and amplified according to the TruSeq^™^ RNA Sample Preparation v2 LS protocol. Finally, the products were purified and enriched with 10 cycles of PCR to create the final doublestranded cDNA library. Final libraries were analyzed using Agilent High Sensitivity DNA analysis chip (Cat. no.5067-4626, Agilent) to estimate the quantity and size distribution, and were then quantified by qPCR using the KAPA Library Quantification Kit (KK4824, KAPA). The prepared library was pooled for single-end sequencing using Illumina HiSeq 2500 with 101-bp single-ended sequence read. Transcript abundance was calculated by first mapping reads to the Arabidopsis TAIR10 genome using Bowtie2 (Langmead and Salzberg, 2012). Unmappable reads were mapped to the TAIR 10 genome sequence by BLAT (Kent, 2002). Read counts were computed using the RackJ package (http://rackj.sourceforge.net/) and normalized using the TMM-quantile method (Robinson and Oshlack, 2010). Normalized read counts were transformed into normalized RPKM values. Z-test was carried out for detecting differentially expressed genes.

Gene ontology (GO) enrichment analysis was analyzed using the Singular Enrichment Analysis (SEA) available on the ArgiGO v2.0 toolkit (Tian et al., 2017). The analysis was performed using the following parameters: selected species: *Arabidopsis thaliana;* Reference: TAIR genome locus (TAIR10_2017); Statistical test method: Fisher; Multi-test adjustment method: Yekutieli (FDR under dependency); Significance level: 0.05; Minimum number of mapping entries: 5; Gene ontology type: Complete GO. Significantly enriched GO terms were summarized and visualized using REVIGO (Supek et al., 2011), with a similarity setting of 0.7 and SimRel as the semantic similarity measure. Final figures were plotted in R (version 3.6.2)

## RESULTS AND DISCUSSION

### Ambient pH Profoundly Alters the Transcriptomic Profile of Iron-Deficient Plants

To investigate the influence of ambient pH on the global gene expression profile of iron-deficient plants, *Arabidopsis* seedlings were grown for 14 days on either optimal (5.5) or high pH (7.0) iron-deficient media, and roots were subjected to transcriptome profiling by RNA-seq. Approximately 24 million reads per treatment were captured during sequencing on the Illumina HiSeq 2000 platform, and mapped to the TAIR10 annotation of the *Arabidopsis* genome.

A flowchart of the experiment is depicted in **Figure 1A**. After filtering lowly expressed genes which may not be of biological relevance (expression level in RPKM < the square root of the mean expression value of the whole dataset), a total of 857 genes were defined as being differentially expressed between iron-deficient plants grown at optimal or high pH media with a < 0.5 or > 2-fold change and a *P* value of < 0.05 (**Fig. 1A**). The expression of a suite of 37 differentially expressed genes (DEGs) was changed more than tenfold (**Supplemental Table 1**), indicative of robust alterations in transcriptional activity by a relatively subtle change in ambient pH.

**Figure 1.**
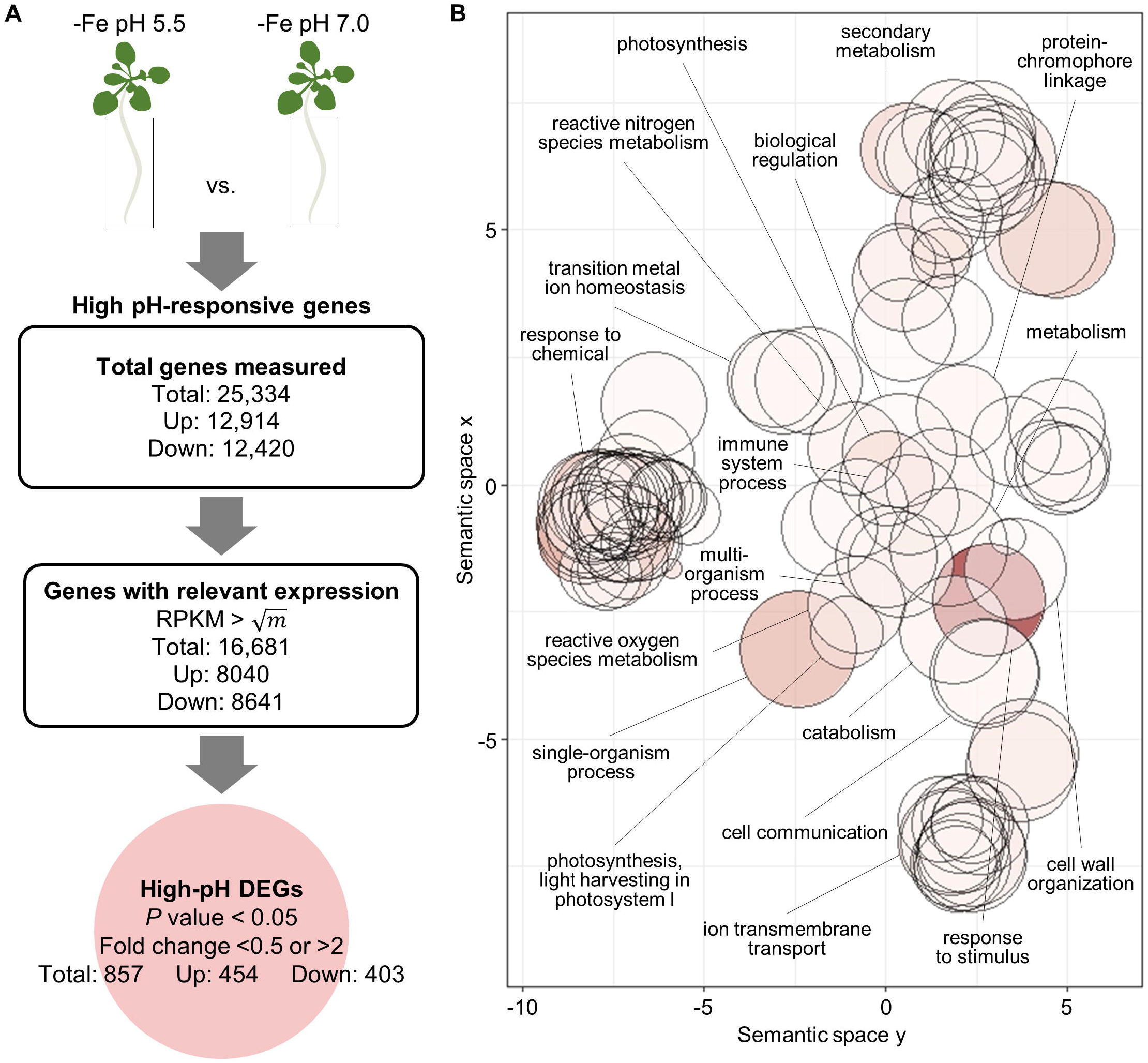
High pH-induced transcriptional changes in iron-deficient plants. A, flowchart of the experiment that depicts the selection criteria used to identify high-pH DEGs. *m* represents the mean of RPKM values in the whole dataset. B, GO biological process term analysis result for the 857 high-pH DEGs as summarized by REVIGO. Bubble color indicates the log10 *P* value of enrichment (bubbles of darker colors are more significant) and the size indicates the frequency of the GO term in the underlying GO annotation database (bubbles of more general terms are larger).

To gain insights into the biological significance of the changes in gene expression, a gene ontology (GO) enrichment analysis for the DEGs was conducted using the Singular Enrichment Analysis (SEA) algorithm available on the AgriGO v2.0 toolkit (Tian et al., 2017), and visualized with the REVIGO Web server (Supek et al., 2011). Categorizing the genes that are responsive to high pH under iron-deficient conditions revealed an overrepresentation of the GO categories ‘response to chemical’ and ‘response to stimulus’, indicating that the majority of pH-responsive genes is involved in adapting the plants to environmental conditions (**Fig. 1B**). Further, the GO category ‘secondary metabolism’ was significantly enriched. Surprisingly, also genes associated with photosynthesis were overrepresented in the data set, probably mirroring photosynthesis-related processes that are mainly affected in leaves where their expression is altered by high pH to avoid photooxidative damage.

Exposure to high pH modulated the iron deficiency response of the plants by altering the expression of several key genes functioning in the acquisition of iron. In high pH plants, *FRO2* transcript levels were reduced by approximately 50% relative to plants grown at pH 5.5 (**Supplemental Table 1**), matching previous observations of severely compromised *in vivo* ferric reduction at high pH (Susin et al., 1996). By contrast, expression of the 2-oxoglutarate-dependent oxygenase *SCOPOLETIN 8-HYDROXYLASE (S8H)* was more than threefold induced when plants were grown on high pH. S8H catalyzes the biosynthesis of the iron-mobilizing coumarin fraxetin (7,8-dihydroxy-6-methoxy-2*H*-chromen-2-one) which is secreted into the rhizosphere under iron-deficient conditions, particularly at high pH (Sisó-Terraza et al., 2016; Siwinska et al., 2018; Tsai and Schmidt, 2018; Rajniak et al., 2018). Fraxetin mobilizes iron by both reduction and chelation, and may form stable complexes with Fe^2+^ (Rajniak et al., 2018; Tsai et al., 2018). Fraxetin-mediated iron reduction is favored by mildly alkaline pH, compensating for the reduced enzymatic ferric reduction activity under such conditions. Interestingly, expression of another iron-responsive gene in the coumarin biosynthesis pathway, the cytochrome P450 *CYP82C4*, was completely abolished when plants were grown on high pH media (**Supplemental Table 1**). CYP82C4 catalyzes the hydroxylation of fraxetin to form sideretin (5,7,8-trihydroxy-6-methoxy-2*H*-chromen-2-one), a catecholic coumarin with a lower pH optimum for iron mobilization when compared to fraxetin (Rajanek et al., 2018). It thus appears that ambient pH can modulate the activity of metabolic enzymes at the transcriptional level to prioritize the production of specific compounds in order to adapt iron acquisition to the prevailing edaphic conditions.

### Ambient pH Modulates the Iron Deficiency Response

A suite of circa 250 genes that are differentially expressed between iron-sufficient and iron-deficient plants surveyed at optimal pH constitute the ‘ferrome’, a robust shift in the transcriptional profile that induces various iron mobilization strategies and aids in recalibrating cellular iron homeostasis (e.g. Rodríguez-Celma et al., 2013; Grillet et al., 2018; Kim et al., 2019). To investigate the effect of a pH shift on iron-deficient plants, a comparative analysis was conducted between high pH-responsive DEGs and DEGs from an RNA-seq-based profiling of iron-sufficient and iron-deficient plants grown under conditions similar to that of the current study except that plants were grown on pH 5.5 media (Rodríguez-Celma et al., 2013). A comparison of the high pH-responsive DEGs with the ferrome revealed an overlap of 54 genes (**Supplemental Table 2**; **Fig. 2a).** Genes from this overlap were mainly enriched in GO terms associated with ‘ion homeostasis’ and ‘response to transition metal nanoparticle’ (**Fig. 2B**). Interestingly, more than half of the genes are oppositely regulated between plants grown at optimal and high pH. Some 74% of the genes that were highly upregulated under iron-deficient conditions at optimal pH were downregulated when plants were grown on high pH media. This group included several key regulators of iron uptake including the subgroup Ib bHLH proteins *bHLH39* and *bHLH101* (Yuan et al., 2008), suggesting a direct or indirect involvement of these proteins in the pH-dependent regulation of iron uptake (**Fig. 2C**). This subset also comprises the iron transporter *IRT2* and the oxidoreductase *FRO2*, which catalyzes the reductive splitting of ferric chelates prior to uptake, the rate-limiting step of iron uptake (Robinson et al., 1999; Grusak et al., 1990; Connolly et al., 2003; Vert et al., 2002).

**Figure 2.**
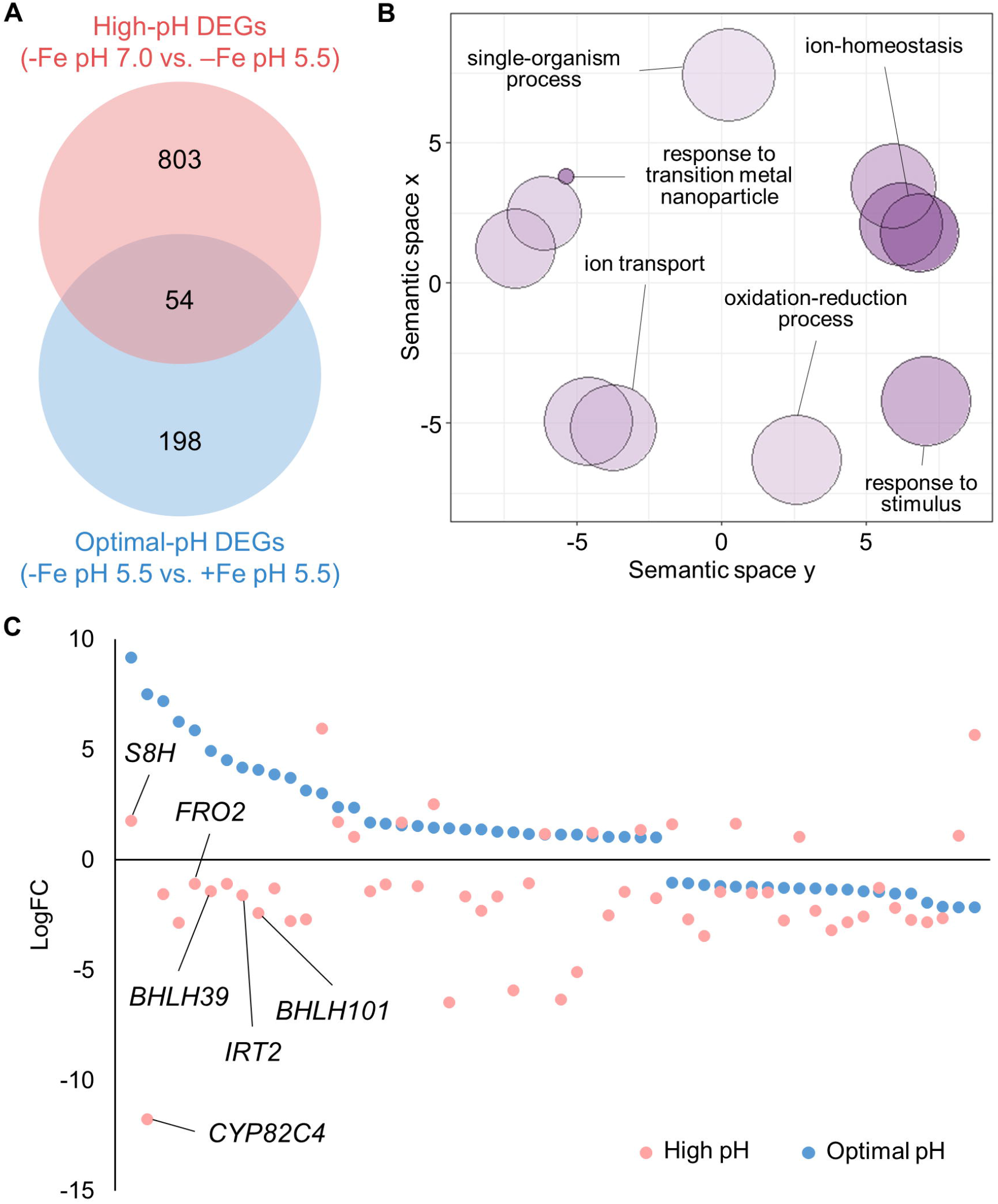
Comparative analysis between high-pH and optimal pH DEGs. A, Venn diagram comparing the DEGs in high-pH and optimal-pH datasets. B, GO biological process term analysis result for the common DEGs in (A) as summarized by REVIGO. Bubble color indicates the log10 *P* value (bubbles of darker color are more significant) and the size indicates the frequency of the GO term in the underlying GO annotation database (bubbles of more general terms are larger). C, Visualization of expression changes of the common DEGs in (A) in both the high-pH and optimal-pH datasets, where the x-axis represents different genes and the y-axis represents the logFC values. Only relevant genes are labeled; refer to Supplemental Table 2 for the complete gene list. Optimal-pH DEGs are from Rodríguez-Celma et al. (2013).

### Exposure of Plants to Different Media pH Uncovers Putative Nodes in pH Signaling

A previously conducted transcriptional survey of *Arabidopsis* roots exposed to low (4.5) pH medium showed that the expression of a total of 1,036 genes was significantly changed after transfer from pH 6.0 (Lager et al., 2010). A comparison of this subset to our high pH data set revealed an overlap of 186 genes. In terms of GO categories, these common DEGs are enriched in the biological functions ‘response to chemical’, ‘immune system process’, ‘cell wall organization and biogenesis’, and ‘regulation of hormone levels’ (**Fig. 3A**). A group of 91 genes were anti-directionally regulated by high and low pH (**Table 1**). The latter list may contain genes that are involved in the perception and transduction of ambient pH. *PLANT INTRACELLULAR RAS GROUP-RELATED LRR 8* (*PIRL8*), a member of a novel class of plant-specific LRR proteins without clearly defined function (Forsthoefel et al., 2005), was found to be highly upregulated in response to high pH and strongly repressed by low pH (**Table 1**). Also, several genes involved in cell wall organization such as the pectin lyase At2g43890, the pectin methylesterase *ATPMEI11* and the expansin *EXP17* showed a large high pH /low pH ratio, possibly to compensate compromised cell expansion at alkaline pH. This supposition is supported by a large high pH/low pH ratio of the auxin-related genes *SAUR41* and *SAUR72*, which have been associated with the regulation of cell expansion (Kong et al., 2013; Qui et al., 2019). High pH /low pH ratios below one were observed for *STRESS INDUCED FACTOR1* (*SIF1*), a root-specific, membrane-anchored LRR kinase with undefined function (Yuan et al., 2018), the aluminum-activated malate transporter *ALMT1*, and the Cys2-His2 zinc-finger domain transcription factor *SENSITIVE TO PROTON RHIZOTOXICITY 2* (*STOP2*). The latter two genes are inducible by low pH stress (Lager et al., 2010; Liang et al., 2013; Kobayashi et al., 2014); almost complete repression was observed under the present (high pH) conditions. Notably, both genes are also induced by phosphate starvation, where ALMT1 and the STOP2 paralog STOP1 repress primary root growth in adaptation to decreased phosphate supply (Balzergue et al., 2017; Wang et al., 2018; Godon et al., 2019). Also, the auxin efflux carrier family proteins *PILS3* and *PILS5* (Barbez et al., 2012) were strongly down- and highly upregulated by high and low pH, respectively, suggesting that ambient pH alters auxin homeostasis. A low high pH/low pH ratio was also observed for *POLYGALACTURONASE INHIBITING PROTEIN 1* (*PGIP1*). PGIP1 stabilizes the cell wall under acidic conditions and was found to be dependent on STOP1/STOP2 (Kobayashi et al., 2014). Notably, *PGIP1*, *STOP2*, *ALMT1*, *SIF1*, and the S-adenosyl-L-methionine-dependent methyltransferases superfamily protein At2g41380 are regulated by STOP1 (Kobayashi et al., 2014).

**Figure 3.**
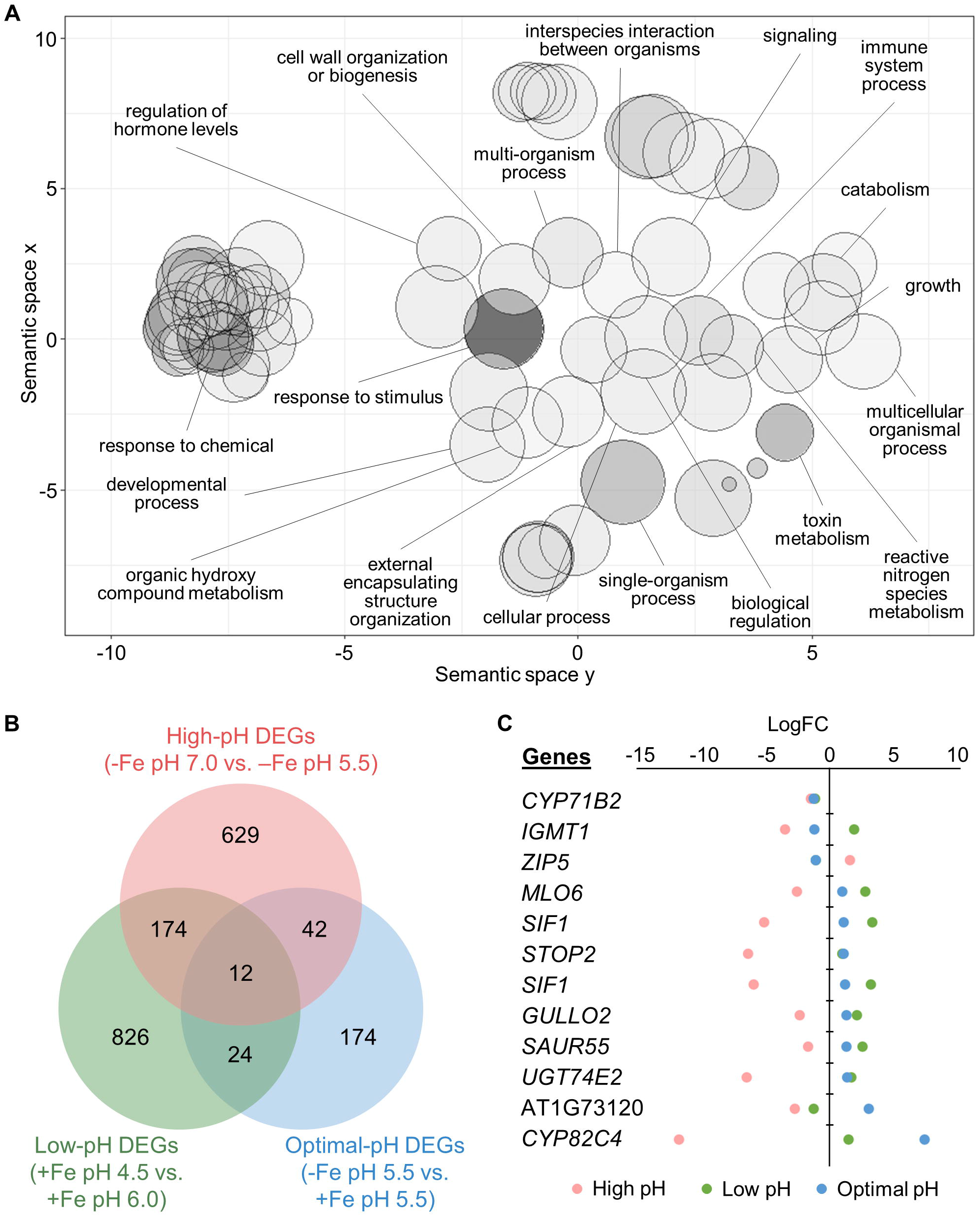
Effect of pH on the transcriptional profile changes in iron-deficient plants. A, GO biological process term analysis result for the 186 common DEGs between high-pH and low-pH datasets as summarized by REVIGO. Bubble color indicates the log10 *P* value (bubbles of darker color are more significant) and the size indicates the frequency of the GO term in the underlying GO annotation database (bubbles of more general terms are larger). B, Venn diagram comparing the DEGs in high-pH, optimal-pH, and low-pH datasets. C, Visualization of expression changes of the 12 common DEGs in (B). The x-axis represents the logFC values and the y-axis represents different genes. Low-pH DEGs are from Lager et al. (2013).

A subset of 12 genes was responsive to all three conditions under investigation and thus represents genes that are highly responsive to both changes in ambient pH and iron supply (**Fig. 3C**). This subset comprises *CYP82C4* and various regulators such as *SIF1*, *STOP2*, and the FIT-regulated F-box/RNI superfamily protein At1g73120. Notably, *CYP82C4*, which was highly upregulated and downregulated under iron deficiency at optimal and high pH, respectively, was also upregulated at low pH conditions even when iron is sufficient, suggesting that the expression of *CYP82C4* is dictated by external pH independent of the iron status.

### Ambient pH is a Major Determinant for the Prioritization of Stress Responses

The transcriptional response of iron-deficient plants grown at high pH also revealed a pronounced overrepresentation of several categories containing genes that orchestrate the defense responses to pathogens, including ERF- and WRKY-type transcription factors, PR proteins, and genes involved in proteolysis, secondary metabolism, and hormone signaling (**Supplemental Figure 1**). All these responses are much less pronounced in iron-deficient plants grown at optimal pH. Exposure to low pH with sufficient iron supply, on the other hand, elicited a more pronounced pathogen response, indicating pH-dependent prioritization of the responses to pathogen attack and iron starvation. In particular, genes related to signaling were overrepresented under low pH conditions. This survey shows that ambient pH considerably modulates the responses to environmental cues by altering the transcriptional landscape of iron-deficient plants to secure and optimize fitness of the plant under a given set of environmental conditions.

## CONCLUSIONS

Alkaline soils are thought to aggravate iron deficiency by rendering the acquisition of iron pools more difficult due to decreased iron activity. Our data show that a difference in ambient pH from 5.5 to 7.0 is causative for considerable differentiation of the global gene expression profile, which goes well beyond simple intensification of the iron deficiency response. A relatively large set of genes is anti-directionally regulated by high and low pH, indicating that ambient pH is translated into transcriptional changes that adapt the plant to the prevailing hydrogen activity, a response that appears to be at least partly elicited by pH *per se* and independent of alterations in the availability of essential minerals such as phosphate and nitrate or toxic elements such as aluminum. Moreover, the iron deficiency response as such is modulated by alterations in ambient pH. While *FRO2* expression is diminished at circumneutral pH, the production and secretion of iron-mobilizing coumarins is induced by high pH, prioritizing the most effective strategy to mobilize iron from otherwise inaccessible pools.

## AUTHOR CONTRIBUTIONS

H.H.T. contributed to the design of the study, carried out experiments, interpreted data and contributed to the manuscript. W.S. conceived and designed the study, participated in the analysis and interpretation of the data and wrote the manuscript. All authors read and approved the final manuscript.

## ACKNOWLEDGEMENTS

We thank the Genomic Technology Core Laboratory at the Institute of Plant and Microbial Biology (IPMB), Academia Sinica, for preparing RNA-seq libraries for sequencing. We also thank Wen-Dar Lin from the Bioinformatics Core Laboratory at IPMB for bioinformatics support.

## SUPPLEMENTAL DATA

**Supplemental Table 1.** High-pH DEGs.

**Supplemental Table 2.** Common DEGs between high-pH and optimal-pH datasets.

**Supplemental Figure 1. MapMan visualization of the biotic stress pathway for the DEGs from different transcriptome datasets.** A, Optimal pH DEGs. B, High pH DEGs. C, 8 h low pH DEGs. Red boxes denote up-regulated genes, blue boxes indicate down-regulated genes. Optimal-pH DEGs are from Rodríguez-Celma et al. (2013). Low-pH DEGs are taken from Lager et al. (2013).

