## Supplemental Figure 1 for "pH-Dependent Transcriptional Profile Changes in Iron-Deficient *Arabidopsis* Roots"

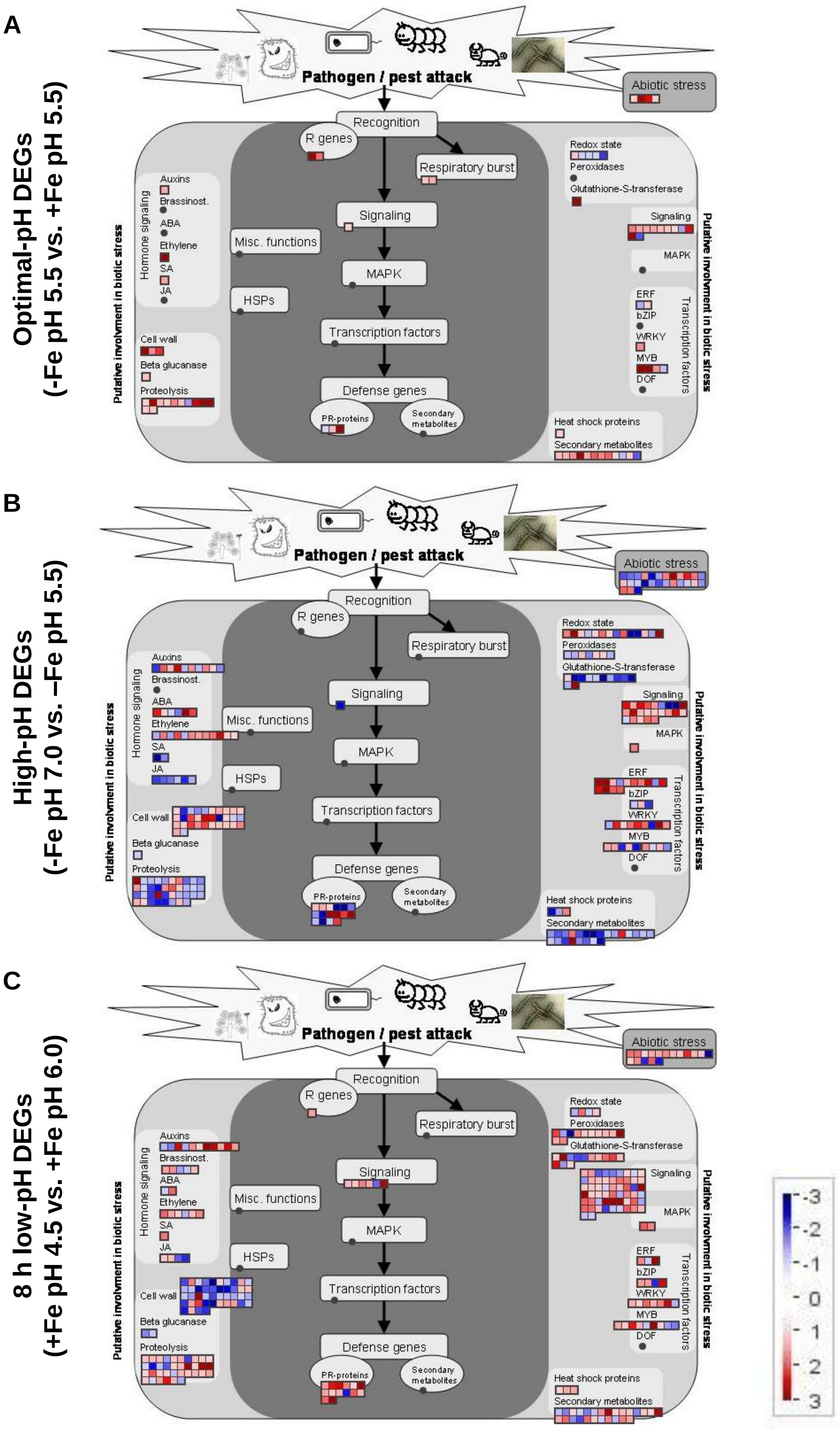


**Supplemental Figure 1. MapMan visualization of the biotic stress pathway for the DEGs from different transcriptome datasets.** A, Optimal pH DEGs. B, High pH DEGs. C, 8 h low pH DEGs. Red boxes denote up-regulated genes and blue boxes denote down-regulated genes. Optimal-pH DEGs are from Rodríguez-Celma et al. (2013). Low-pH DEGs are from Lager et al. (2013).
